# Environmental DNA concentrations of Japanese eels in relation to habitat characteristics

**DOI:** 10.1101/2022.09.29.510209

**Authors:** Yurika Ono, Katsuya Hirasaka, Taijun Myosho, Shingo Fujimoto, Mitsuharu Yagi

**Affiliations:** Graduate School of Fisheries and Environmental Sciences, Nagasaki University, 1-14 Bunkyo, Nagasaki 852-8521, Japan; Organization for Marine Science and Technology, Nagasaki University, Nagasaki 852-8521, Japan; Graduate Division of Nutritional and Environmental Sciences, University of Shizuoka, Shizuoka 422-8526, Japan; Tropical Biosphere Research Center, University of the Ryukyus, Okinawa 905-0227, Japan; Institute of Integrated Science and Technology, Nagasaki University, 1-14 Bunkyo, Nagasaki 852-8521, Japan

**Author notes:** Corresponding author; E-mail, (M. Yagi).

**Keywords:** *Anguilla japonica*, eDNA, distribution, Nagasaki

## Abstract

The Japanese eel (*Anguilla japonica*), is listed as “Endangered” by the IUCN. Understanding eel riverine habitat is useful in considering conservation strategies. This study sought to determine the relationship between environmental DNA (eDNA) concentrations derived from Japanese eels, water quality, and river structure in three small rivers in Nagasaki, Japan. eDNA was detected at 14 of 15 sites (93%). The concentration of eDNA in brackish water was significantly higher than that in freshwater and was correlated with water depth. Eel occurrence throughout the river suggests a need to conserve a diversity of habitats.

---

Populations of *Anguilla* spp. are shrinking worldwide, and conservation, as well as habitat monitoring of this genus, is an urgent issue (Itakura et al., 2020). There are 16 species in the genus *Anguilla* worldwide, distributed from temperate to tropical zones. Populations of the genus *Anguilla* occur in more than 150 countries and are of ecological, commercial, and cultural importance (Jacoby et al. 2015). However, in recent years, eels have declined significantly, and 10 species, including the Japanese eel (*Anguilla japonica*), are listed as “Endangered” on the IUCN Red List of Threatened Species (IUCN 2022).

Riverine habitats are essential for maturation of Japanese eels (Kaifu et al. 2010). The Japanese eel is migratory fish that spawns in the open ocean west of the Mariana Islands (Aida et al. 2003; Tsukamoto et al. 2011). After hatching, leptocephalus larvae are transported westward by the North Equatorial Current and then northward by the Kuroshio Current, approximately six months before they reach coastal areas of East Asia (Kimura et al. 1994). Leptocephalus larvae approach coastal areas, metamorphose into glass eels and migrate along coasts, where they ascend rivers (Arai et al. 2003). Eels shift to a benthic lifestyle near the estuary and then slowly grow and spread upstream or downstream, including coastal areas (Moriarty 2003; Kaifu 2016).

Although *Anguilla japonica* occurs at low densities, distributional information is vital for its conservation (Fukukmoto et al. 2015; Sakata et al. 2017). In recent years, analytical methods to monitor distribution of populations using environmental DNA (eDNA) have seen increasing use (Lodge et al. 2012). eDNA is a generic term for DNA present in the environment, which is derived from excrement, mucus, blood, and sloughed cells (Minamoto et al. 2016). This analytical method has the advantage that it can be used safely and without contact with target organisms, and that a large area can be surveyed easily and at low cost (Jerdge et al. 2011; Takahara et al. 2012). It is also more sensitive for detecting fish species than traditionally used monitoring methods, such as fishing, nets, electric shockers and observation (Jerde et al. 2013; Itakura et al. 2019; 2020). A study on the Japanese eel using eDNA analysis reported high environmental DNA concentrations along the west coast of Kyushu, the Pacific side of Honshu and the Seto Inland Sea, accounting for 80% of all sites across Japan (Kasai et al. 2021). Itakura et al. (2019) showed that eDNA analysis can be used to estimate populations and biomass of Japanese eels in rivers. Eels are thought to migrate and grow in brackish and freshwater areas as they search for suitable habitats, but there are few studies examining the relationship between the physical river environment and eDNA concentrations. Thus, this study sought to determine the relationship between Japanese eel environmental DNA concentrations, water quality, and river structure in several small rivers.

In the present study, eDNA surveys were conducted with a focus on yellow eels (*Anguilla japonica*). Yellow eels spend several years to decades growing in rivers, a relatively sedentary period in the life history of this species (Itakura et al. 2019.). As the glass eel upwelling and silver eel migration occur in autumn and winter (Itakura et al. 2019; Sudo et al. 2017), eDNA surveys in this study were carried out between July and September 2021 to avoid collecting eDNA derived from glass eels. Water sampling was conducted in Nagasaki Prefecture, Kyushu, Japan, in the Nakajima River, the Emukae River and the Arie River (Fig. 1a). These three are small rivers, less than 20 km long and with a collective basin area of 100 km^2^. Water sampling was carried out at 4-6 sites per river, for 15 sites from the lower to the upper reaches of these rivers (Fig. 1 and 2). At sampling stations, water temperature, dissolved oxygen, salinity, and depth were measured using a multiparameter digital water quality meter (YSI ProDSS, Xylem Inc.). In addition, flow velocity, elevation, river width, fish and other organisms within visual inspection were recorded.

**Fig. 1.**
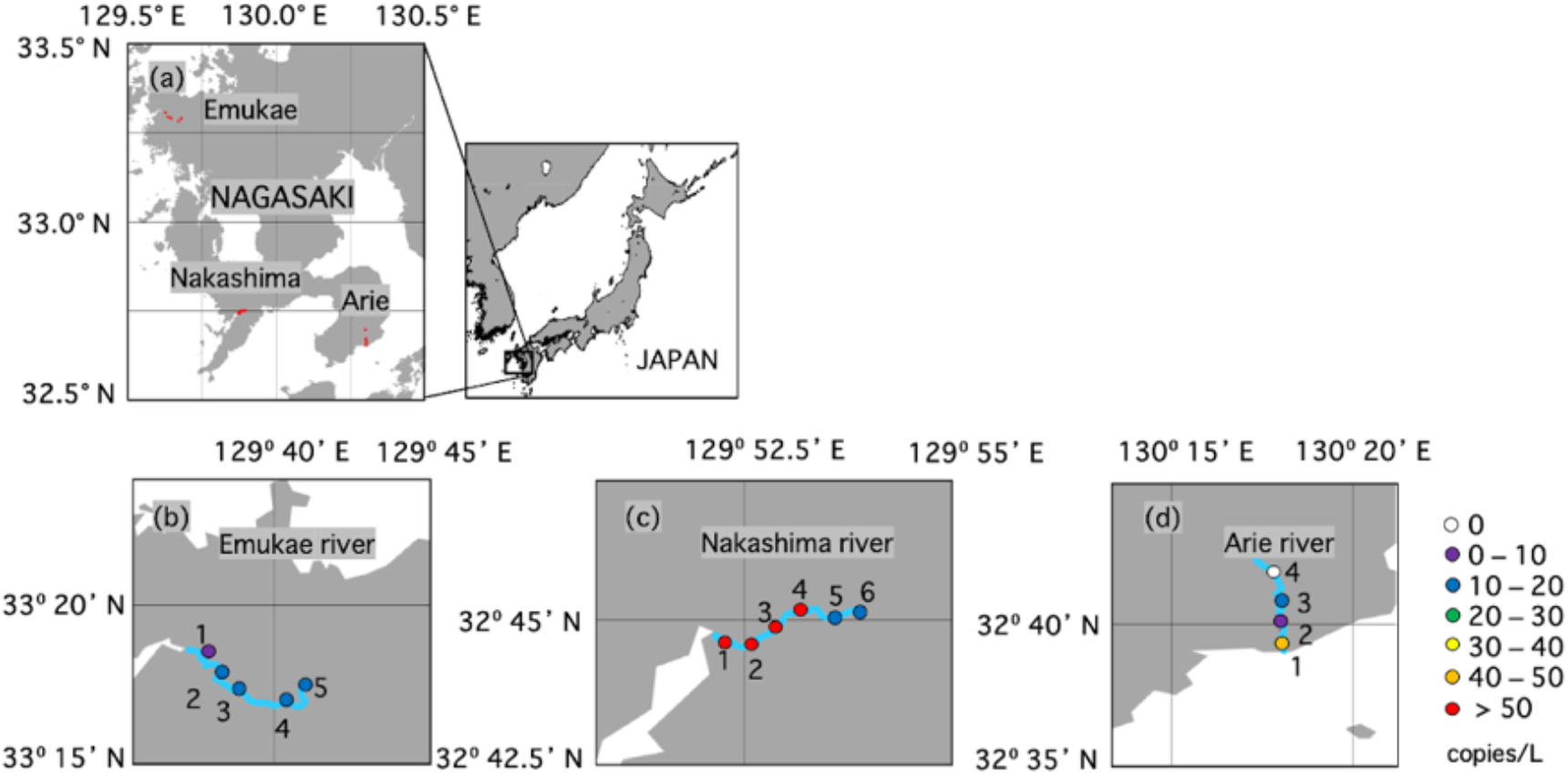
Map showing survey sites and environmental DNA concentrations of the Japanese eel, *Anguilla japonica*. (a) Geographical location of study sites in Kyushu, Japan, (b) Emukae River, (c) Nakashima River, (d) Arie River. Solid circles indicate sites where water samples were collected.

**Fig. 2.**
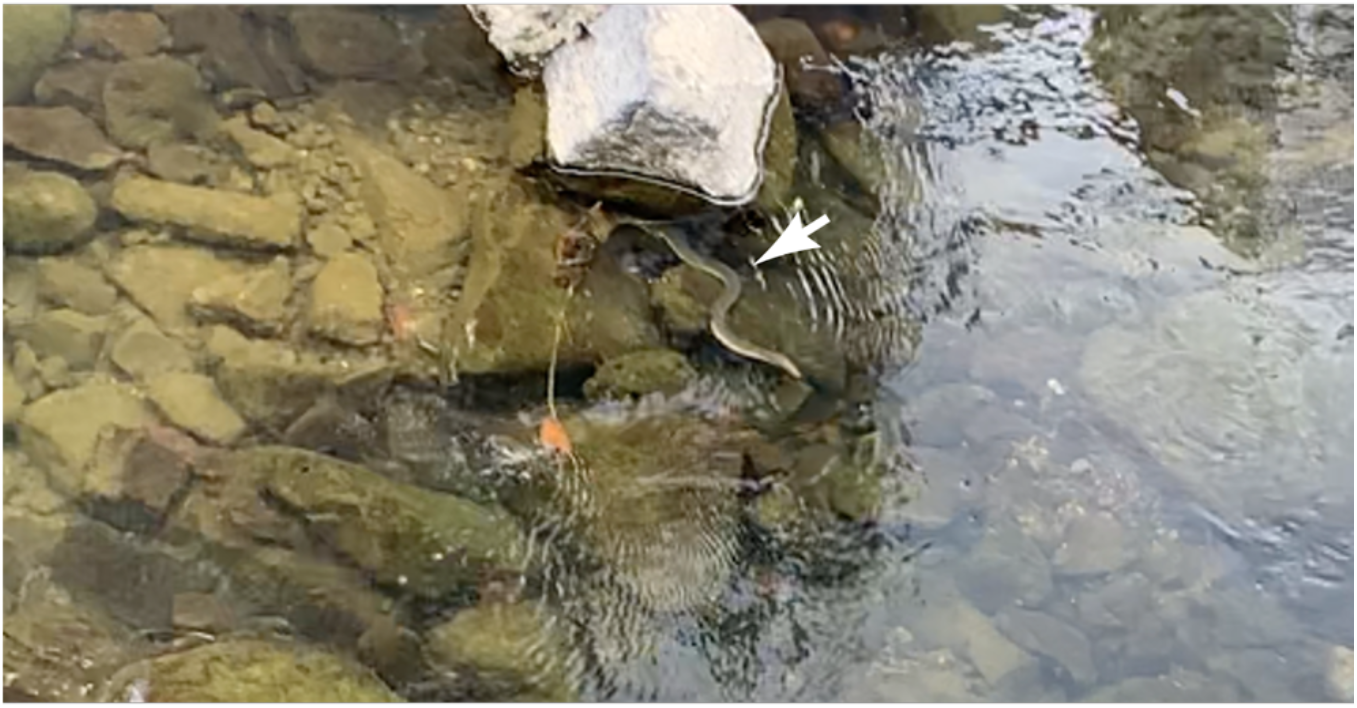
Swimming Japanese eel observed during water sampling at Site 4 on the Nakashima River,
Nagasaki, Japan.

Water sampling was carried out according to methods in the manual for aquatic environment DNA studies and experiments (Minamoto et al. 2020). After collecting 1 L of water using a disposable cup, 1 mL of 10% benzalkonium chloride solution (Final concentration of 0.01% v/v) was immediately added to the sample to inhibit eDNA degradation (Yamanaka et al. 2017). At each site, water sampling was conducted in main channel of the river, and at sites where the main channel was inaccessible, water sampling was carried out from a bridge using a fishing rod and reel. Water sampling was performed by the same investigator throughout the study. To avoid contamination from other sites, powder-free nitrile gloves were worn during water sampling and were changed at each site. After sampling, water was transported in a cooler with ice to the Fish and Ships Laboratory at Nagasaki University, Japan. Bottles containing 1 L of distilled water and 1 mL of 10% benzalkonium chloride solution were also placed in the cooler box and used as blank controls. Blanks were prepared for each sample. Samples and blanks were vacuum filtered through one or two 47 mm GF/F glass filters (pore size: 0.7 μm, GE Healthcare, Tokyo, Japan) within 48 h. Filters were wrapped in aluminum foil and stored at −20 °C until eDNA extraction.

eDNA extraction followed the method of Itakura et al. (2019). Briefly, total DNA was extracted from each filter using a DNeasy Blood and Tissue Kit (Qiagen, Hilden, Germany). DNA recovery was performed by a spin column method according to the manual provided. Blanks were processed using the same process. Recovered DNA, as well as blanks, were stored at −20 °C until the Applied Biosystems 7500 Fast real-time PCR assay (qPCR).

eDNA assays were performed according to the protocol of Itakura et al. (2019) using a dilution series of synthetic linear DNA (231bp: 5’-ATACACTCCCCACCCCCTAAAAATATTAAGCTATCCTATGCACACATAGGAGAAACAATGCTAAAATCAGTAATAAGAGGGCCCAAGCCCTCTCCTAGCACATGTGTAAGTCAGAACGGACCGACCACTGACAATCAACGGACCCAAACAGAGAGAAAAAGAACAAACTACAAAAAACAAGAAAAATCTATTTAATACCACAAACCGTTAACCCAACACAGGAGTGCCTAA-3’) as a quantitative standard (dilutions of 1×10^1^, 1×10^2^, 1×10^3^, 1×10^6^ and 1×10^9^) for all qPCR assays. In each round of qPCR, three replicates per sample and pure water (2 μL) as a negative control were analysed. In all rounds, qPCR calibration curves had *R^2^* values ranging from 0.997 to 0.998, slopes from −3.641 to −3.596 and intercepts from 39.426 to 42.468. Copy numbers were calculated based on calibration curves for each round and Ct values for each sample.

Preliminary experiments using water from Japanese eel rearing tanks confirmed that eel eDNA could be detected. One Japanese eel (total length: 49 cm, wet weight: 137.9 g) was collected with a dip net in the Urakami River, Nagasaki, Japan. Eel was reared in 55 L tanks at a water temperature of 20 °C. After approximately 1 month of acclimatisation, 1 L of rearing water was sampled and dilution series of 1×10^−1^, 1×10^−2^, 1×10^−3^, 1×10^−4^, 1×10^−5^, 1×10^−6^ and 1×10^−7^ were made using distilled water. Three replicates of sample water were prepared, each with 1 mL 10% benzalkonium chloride. Extraction and quantification of eDNA were carried out by the methods described above. The results showed that eel eDNA was detected, with mean copy numbers/L ± s.d. (n) of 7290 ± 4876 (3), 753 ± 393 (3), 18.7 ± 12.3 (3), 4.52 (1), 8.18 (1) at 1, 10^−1^, 10^−2^, 10^−3^ and 10^−4^-fold, respectively, and not detectable at 10^−5^, 10^−6^ and 10^−7^. This experiment was approved by the Animal Care and Use Committee of the Faculty of Fisheries, Nagasaki University (permission no. NF-0055), in accordance with Guidelines for Animal Experimentation of the Faculty of Fisheries (fish, amphibians, and invertebrates), Nagasaki University.

Statistical analysis was performed by pooling data from all three rivers. Brackish and freshwater areas were divided on the 5‰ criterion based on salinity as measured by water quality (Lai et al., 2015). Data were checked for normality using the Shapiro-Wilk test. Student t-tests were conducted to determine whether differences in mean eDNA concentrations between brackish and freshwater areas were significant. Pearson’s correlation coefficient test examined the relationship between eDNA concentration and each physical environmental parameter. The significance level was set at *p* = 0.05 and the statistical software JMP Pro 16 was used.

Eel-derived eDNA was detected in all three rivers and at 14 of the 15 sites (Fig. 2). It was not detected at the uppermost site on the Ariake River (Fig. 2c). Itakura et al. (2019) reported that eDNA was detected in 100% (27 sites) of the three rivers in Kyushu, Japan. The detection rate was over 80% in western Kyushu, the Pacific side of the main island of Japan, and the Seto Inland Sea (Kasai et al. 2021). In this study, the detection rate at sites from downstream to upstream was 93%, which is similar to the former studies, indicating that Japanese eels inhabit almost all parts of the river. In fact, direct capture methods have confirmed that Japanese eels inhabit a wide range of habitats, from clear streams and rapids in the upper reaches of rivers and downstream, to highly polluted waters (Itakura et al. 2019). It will be important for eel conservation to maintain continuity in riverine and coastal areas and to conserve and create diverse riverine habitats.

eDNA copy numbers were highest at Site 3 on the Nakashima River at 96.1 copies/L and lowest at Site 1 on the Emukae River at 6.2 copies/L. Swimming yellow eels were observed at S4 of the Nakajima River, approximately 10 m upstream of the sampling point (Fig. 3), and this site was 92.4 copies/L (Fig. 2). eDNA concentrations of Japanese eels in rivers in Japan reported to date vary considerably. While Kasai et al. (2021) reported that 44356 copies/L were observed in the Asaragi River on the Pacific side of the main island, the maximum was approximately 250 copies/L in the Atsumari river in Kyushu. In preliminary experiments for this study, the concentration in the rearing water in a 55 L tank was 7290 copies/L, which means that 50 copies/L is the concentration of an adult eel in approximately 8 tonnes of water. In fact, analyses based on direct capture methods have reported a weak, but significant correlation between eel biomass and abundance with eDNA concentrations (Itakura et al. 2019). Although this experiment is limited by the lack of direct capture, eDNA concentrations in this study are also likely to reflect the biomass and abundance of yellow eels.

**Fig. 3.**
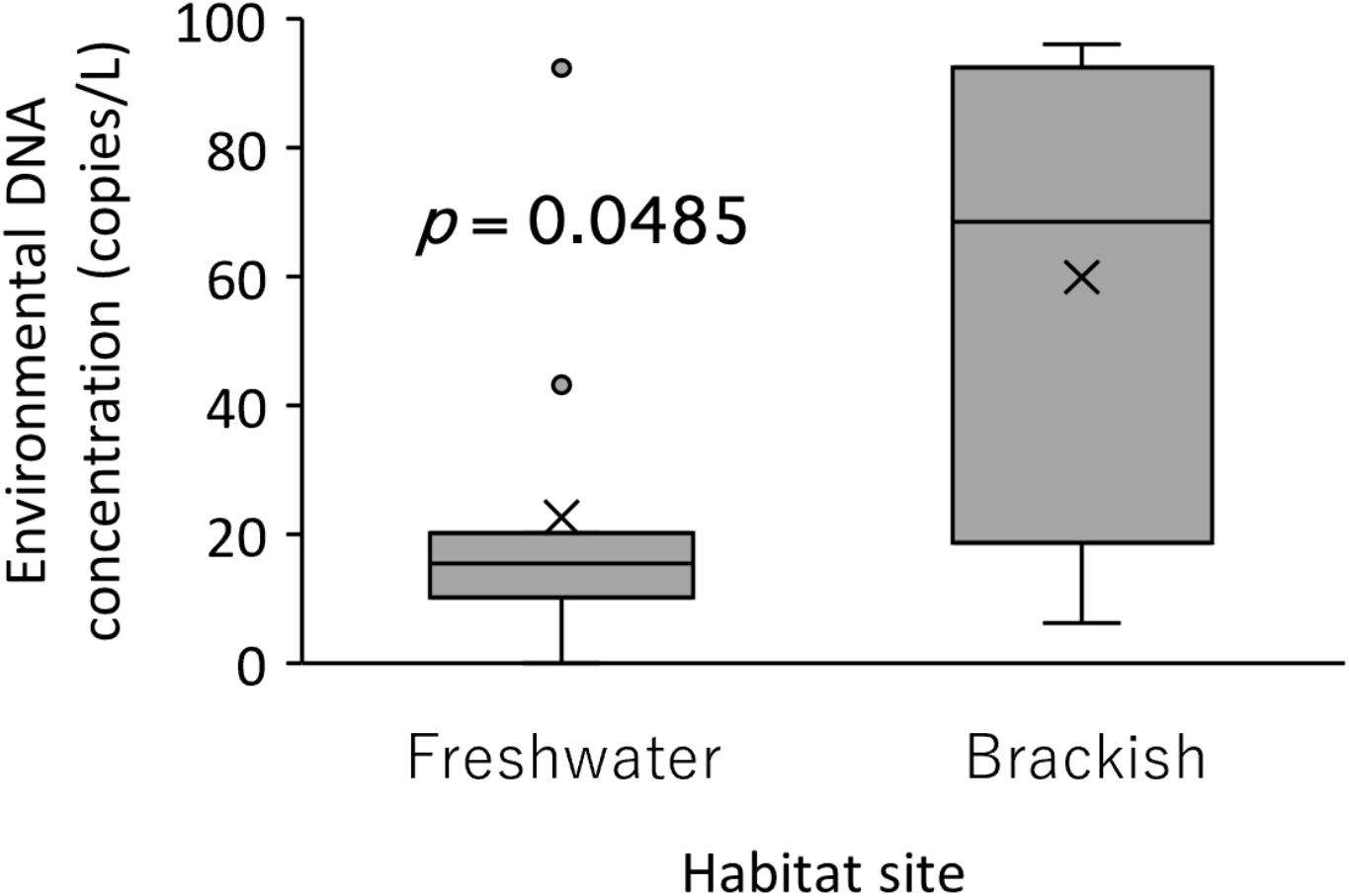
Comparison of environmental DNA concentration of Japanese eels in brackish and freshwater.

eDNA concentrations were 59.9 ± 34.2 copies/L (mean ± S.D) (n = 4) in brackish water and 22.6 ± 24.3 (n = 11) in freshwater. This difference was significant (*p* < 0.05) (Fig. 3). This result confirmed the finding of Itakura et al. (2020), showing that eDNA concentrations are higher in downstream than upstream sites. Previous studies also showed that the density of eels decreases with increasing distance upstream from the river mouth (Yokouchi et al. 2008; Kume et al. 2020). In this study, we also found that there was a significant correlation between eDNA concentration and depth (*R* = 0.60, *p* < 0.05). However, water temperature, dissolved oxygen, salinity, flow velocity and elevation showed no significant correlation with eDNA concentration, implying that eels inhabit a wide variety of aquatic environments. Further studies on the physical environment as well as on sympatry with river structures and other organisms are needed to understand eel habitats.

Japanese eels have been stocked in rivers throughout Japan for population restoration (Kaifu et al. 2019). However, according to the Fishery Census 2008, 2013, no releases have been conducted in Nagasaki Prefecture. Therefore, eDNA detected in this study is most likely to be from naturally occurring eels. Documenting the distributions and biomasses of Japanese eels in Nagasaki Prefecture and comparing them with rivers where releases have been carried out may provide useful information to determine the ecological consequences and effectiveness of releases.

## Acknowledgements

This work was supported by JSPS KAKENHI (Grant Number JP21K06337 to MY). We are grateful to students from the “Fish and Ships Laboratory” in the Faculty of Fisheries, Nagasaki University, who assisted us with the study. Finally, thanks to the editors and anonymous reviewers for their valuable comments and suggestions that greatly improved the quality of the manuscript.

## References

Fukuoka, A., T. Takahara and M. Matsumoto (2016) Establishment of detection system for native rare species, *Hemigrammocypris rasborella*, using environmental DNA. Jpn. J. Ecol, 66, 613–620 (in Japanese with English abstract).

Fukumoto, S., A. Ushimaru and T. Minamoto (2015) A basin-scale application of environmental DNA assessment for rare endemic species and closely related exotic species in rivers: A case study of giant salamanders in Japan. J. Appl. Ecol., 52, 358–365.

Itakura H., R. Wakaki., M. Gollock and K. Kaifu (2020) Anguillid eels as a surrogate species for conservation of freshwater biodiversity in Japan. Sci. Rep., 10, 8790

Itakura, H., R. Wakiya., M. K. Sakata., H. Hsiang-Yi., C. Shih-Chong., Y. Chih-Chao., H. Yi-Cheng., H. Yu-San., S. Yamamoto and T. Minamoto (2020) Estimations of Riverine Distribution, Abundance, and Biomass of Anguillid Eels in Japan and Taiwan Using Environmental DNA Analysis. Zool. Stud., 59, e17.

Itakura, H., R. Wakiya., S. Yamamoto., K. Kaifu., T. Sato and T. Minamoto (2019) Environmental DNA analysis reveals the spatial distribution, abundance, and biomass of Japanese eels at the river-basin scale. Aquatic. Conserv. Mar. Freshw. Ecosyst., 29,361–373.

International Union for the Conservation of Nature (IUCN) (2022) The IUCN red list of threatened species. Version 2022.3

Jacoby, D. M. P., J. M. Casselman., V. Crook., M.-B. DeLucia., H. Ahn., K. Kaifu., T. Kurwie., P. Sasal., A. M. C. Silfvergrip., K. G. Smith., K. Uchida., A. M. Walker and M. J. Gollock (2015) Synergistic patterns of threat and the challenges facing global anguillid eel conservation. Glob. Ecol. Conserv., 4, 321–333.

Jerde, C. L., W. L. Chadderton., A. R. Mahon., M. A. Renshaw., J. Corush., M. L. Budny., S. Mysorekar and D. M. Lodge (2013) Detection of Asian carp DNA as part of a Great Lakes basin-wide surveillance program. Can. J. Fish. Aquat. Sci., 70, 522–526

Jerde, C. L., A. R. Mahon., W. L. Chadderton and D. M. Lodge (2011) “Sight-unseen” detection of rare aquatic species using environmental DNA. Conserv. Lett., 4, 150–157

Yokouchi, K., Y. Kaneko., K. Kaifu., J. Aoyama., K. Uchida and K. Tsukamoto (2014) Demographic survey of the yellow-phase Japanese eel Anguilla japonica in Japan. Fish. Sci., 80, 543–554.

Kaifu, K (2019) Challenges in assessments of Japanese eel stock. Mar. Policy., 102, 1–4.

Kaifu, K., M. Tamura., J. Aoyama and K. Tsukamoto (2010) Dispersal of yel low phase Japanese eels Anguilla japonica after recruitment in the Kojima Bay-Asahi river system, Japan. Environ. Biol. Fishes, 88, 273–282.

Kasai, A., A. Yamazaki., H. Ahn., H. Yamanaka., S. Kameyama., R. Masuda., N. Azuma., S. Kimura., T. Karaki., Y. Kurokawa and Y. Yamashita (2021) Distribution of Japanese Eel Anguilla japonica Revealed by Environmental DNA. Front. Ecol. Evol., 9, 621461

Kimura, S., K. Tsukamoto and T. Sugimoto (1994) A model for the larval migration of the Japanese eel: roles of the trade winds and salinity front. Mar. Biol., 119(2), 185–190

Kume, M., Y. Terashima., F. Kawai., A. Kutzer., T. Wada and Y. Yamashita (2020) Size-dependent changes in habitat use of Japanese eel Anguilla japonica during the river life stage. Environ. Biol. Fishes, 103, 269–281.

Lai, Z., R. Ma., G. Gao., C. Chen and R. C. Beardsley (2015) Impact of multichannel river network on the plume dynamics in the Pearl River estuary. J. Geophys. Res. Ocean. 120, 5766–5789.

Lodge, D. M., C. R. Turner., C. L. Jerde., M. A. Barnes., L. Chadderton., S. P. Egan and M. E. Pfrender (2012) Conservation in a cup of water: Estimating biodiversity and population abundance from environmental DNA.Mol. Ecol., 21, 2555–2558.

Sakata, M. K., T. Watanabe., N. Maki., K. Ikeda., T. Kosuge., H. Okada., T. Sado., M. Miya and T. Minamoto (2021) Determining an effective sampling method for eDNA metabarcoding: a case study for fish biodiversity monitoring in a small, natural river. Limnology, 22, 221–235.

Minamoto, T., T. Naka., K. Moji and A. Maruyama (2016) Techniques for the practical collection of environmental DNA: filter selection, preservation, and extraction. Limnology, 17, 23–32

Minamoto, T., M. Miya., T. Sado., S. Seino., H. Doi and M. Kondoh (2020) An illustrated manual for environmental DNA research: water sampling guidelines and experimental protocols. Environ. DNA, 3, 8–13.

Minamoto, T., M. Miya., T. Sado., S. Seino., H. Doi., M. Kondoh., K. Nakamura., T. Takahara., S. Yamamoto., H. Yamanaka., H. Araki., W. Iwasaki., A. Kasai., R. Masuda and K. Uchii (2021) An illustrated manual for environmental DNA research: Water sampling guidelines and experimental protocols. Environ. DNA, 3, 8–13.

Moriarty C (2003) The yellow eel. In “Eel Biology” (ed. by K. Aida, K. Tsukamoto and K. Yamauchi), Springer-Verlag, Tokyo, pp. 89–105, 121–140.

Sakata, M. K., N. Maki., H. Sugiyama and T. Minamoto (2017) Identifying a breeding habitat of a critically endangered fish, Acheilognathus typus, in a natural river in Japan. Sci. Nat., 104, 100.

Sudo R., A. Okamura., M. J. Miller and K. Tsukamoto (2017) Environmental factors affecting of spawing migrations of the Japanese eels (*Anguilla japonica*) in Mikawa Bay Japan. Environ. Biol. Fishes, 100, 237–249.

Takahara T, T. Minamoto., H. Yamanaka., H. Doi and Z. Kawabata (2012) Estimation of fish biomass using environmental DNA. PLoS ONE, 7, e35868.

Arai, T., A. Kotake., M. Ohji., N. Miyazaki and K. Tsukamoto (2003) Migratory history and habitat use of Japanese eel *Anguilla japonica* in the Sanriku Coast of Japan. Fish. Sci., 69, 813–818.

Takahara, T., H. Yamanaka., T. Minamoto., H. Doi and K. Uchii (2016) Current state of biomonitoring method using environment DNA analysis. Japanese J. Ecol., 66, 583–600.

Shiragaki, T., T. Inoue., H. Fukuda., M. Ushio., M. Kusaka., T. Okano and H. Takasu (2021) Pilot Study on Fish Species Composition Using Environmental DNA in Marine Sediment. JSWE, 44, 79–84 (in Japanese with English abstract).

Tsukamoto, K., S. Chow., T. Otake., H. Kurogi., N. Mochioka., M. J. Miller., J. Aoyama., S. Kimura., S. Watanabe., T. Yoshinaga., A. Shinoda., M. Kuroki., M. Oya., T. Watanabe., K. Hata., S. Ijiri., Y. Kazeto., K. Nomura and H. Tanaka (2011) Oceanic spawning ecology of freshwater eels in the western North Pacific. Nat. Commun., 2, 179.

Yamanaka, H., T. Minamoto., J. Matsuura., S. Sakurai., S. Tsuji., H. Motozawa., M. Hongo., Y. Sogo., N. Kakimi., I. Teramura., M. Sugita., M. Baba and A. Kondo (2017) A simple method for preserving environmental DNA in water samples at ambient temperature by addition of cationic surfactant. Limnology, 18, 233–241.

Yokouchi, K., J. Aoyama., H. P. Oka and K. Tsukamoto (2008) Variation in the demographic characteristics of yellow-phase Japanese eels in different habitats of the Hamana Lake system, Japan. Ecol. Freshw. Fish., 17, 639–652.

